# Nuclear gene proximity and protein interactions shape transcript covariances in mammalian single cells

**DOI:** 10.1101/771402

**Authors:** Marcel Tarbier, Sebastian D. Mackowiak, João Frade, Silvina Catuara-Solarz, Inna Biryukova, Eleni Gelali, Diego Bárcena Menéndez, Luis Zapata, Stephan Ossowski, Magda Bienko, Caroline J. Gallant, Marc R. Friedländer

## Abstract

Single-cell RNA sequencing studies into gene co-expression patterns could yield important new regulatory and functional insights, but have so far been limited by the confounding effects of cell differentiation and the cell cycle. We apply a tailored experimental design that eliminates these confounders, and report >80,000 intrinsically covarying gene pairs in mouse embryonic stem cells. These covariances form a network with biological properties, outlining known and novel gene interactions. We provide the first evidence that miRNAs naturally induce transcriptome-wide covariances, and compare the relative importance of nuclear organization, transcriptional and post-transcriptional regulation in defining covariances. We find that nuclear organization has the greatest impact, and that genes encoding for physically interacting proteins specifically tend to covary, suggesting importance for protein complex stoichiometry. Our results lend support to the concept of post-transcriptional *‘RNA operons’*, but we further present evidence that nuclear proximity of genes on the same or even distinct chromosomes also provides substantial functional regulation in mammalian single cells.

## Introduction

Two genes that increase or decrease coordinately in expression over multiple conditions are said to covary in expression. Gene expression covariance can be studied over two conditions (for instance healthy and diseased tissue), over time-series experiments, or in meta-studies spanning hundreds of tissues and cell types, for instance from public expression repositories ^1–3^. Over the last 20 years, these studies have yielded numerous important biological insights since covarying genes are often functionally related and commonly share the same gene regulatory mechanisms.

In the last ten years, new single-cell sequencing methods have emerged, making it possible to profile entire transcriptomes of individual cells ^4–6^. This makes it possible to identify genes that covary in expression across individual cells, considering in effect every cell as a distinct condition. This research direction holds great promise, since it could reveal biological covariances that are not detectable in analysis of bulk cell populations. First, differences in cellular compositions between samples may disturb covariance analyses in bulk tissues^7^. Further, transcripts can appear to be constantly and moderately expressed in all studied tissues or cell cultures, but may in fact display temporally fluctuating and covarying expression in single cells. This latter type of covariances may never be detected in bulk tissues. However, until now transcriptome-wide single-cell studies of such intrinsic gene covariance patterns have been limited by confounding factors such as cell cycle progression and cell differentiation, that are extrinsic to the genes of interest^8,9^. These confounding factors induce a strong impact on the global covariance patterns, which could overshadow the more subtle - and potentially more interesting - underlying patterns.

Here, we apply carefully designed experimental conditions to remove the confounding extrinsic effects of differentiation and cell cycle progression, and apply sensitive Smart-Seq2 single-cell sequencing to profile the transcriptomes of hundreds of mouse embryonic stem cells (mESCs). Specifically, using stringent cut-offs we report >80,000 gene pairs that intrinsically covary in expression; more than have been described in previous single-cell studies. The covarying gene pairs interlink to form a network with well-established biological features, following a so-called power-law distribution, and recover known regulatory patterns and pathways. We further apply a novel computational framework to study the relative importance of distinct regulatory mechanisms for gene expression covariances. We find that genes regulated by the same transcription factors or miRNAs tend to covary, while the strongest effect is seen with genes that are in nuclear proximity on the same chromosomes. A similar but weaker effect is seen for genes that are in nuclear proximity on distinct chromosomes. We validate that a subset of the covariances are directly induced by miRNAs by repeating our entire experiment in miRNA-deficient cells.

Last, we test two competing hypotheses regarding the putative function of gene expression covariances. The first hypothesis states that genes covary in expression to ensure stoichiometric abundances of proteins that function in the same pathway, while the second hypothesis proposes that covariances are important for proper stoichiometry of proteins that are part of the same complexes. We find that covarying genes only tend to share the same function if their encoded proteins also physically interact, lending evidence to the “protein complex” hypothesis.

In summary, we have combined single-cell RNA sequencing with a tailored experimental design and novel computational analyses to quantify regulatory drivers in single mammalian embryonic stem cells, highlighting the importance of nuclear proximity for gene expression covariances. Additionally, we present evidence that these covariances play roles in ensuring stoichiometry between interacting proteins.

## Results

### Smart-Seq2 sequencing of hundreds of mouse single-cell transcriptomes

To obtain reliable and reproducible measurements of gene expression for our study, we applied the Smart-Seq2 protocol to sequence the transcriptomes of 567 individual mouse embryonic stem cells divided between three well-plates which serve as biological replicates. The Smart-Seq2 protocol is one of the gold standards in the single-cell field. While labor-intensive and not easily scalable, it is highly sensitive and precise, which means that each measurement is less affected by technical noise^6,10,11^. It also reliably detects both exons and introns, which is useful for distinguishing transcriptional and post-transcriptional regulation^12^. We performed strict quality filtering on the initial set of cells, which resulted in a total of 355 cells considered (Methods, Suppl. Fig. 1, Suppl. Table 1). Gene expression values in each cell were normalized to the sum of mRNA sequence reads in the given cell, as this approach was found to best eliminate biases from technical factors (Methods, Suppl. Fig. 2, 3). We considered a gene to be reliably profiled if its transcript was detected in at least half of the cells in each of the three replicates. Importantly, we estimated that for genes that are expressed at above 16 reads per million biological variation between cells exceeds technical noise (as estimated by ERCC spike-ins, Suppl. Fig. 4). The vast majority of the genes that fulfill our criteria for reliable detection also exceeds this expression threshold. Overall our analysis yielded reliable gene expression measurements for 8,983 genes (Suppl. Table 2).

### Homogenous cell population unconfounded by cell cycle or differentiation

For the sequencing experiment, we took several precautions to eliminate the confounding extrinsic effects of cell cycle and differentiation. First, all cells were cultured in 2i+LIF medium which is a well-established protocol to maintain embryonic stem cells in a homogeneous pluripotent state, excluding potential differentiation effects^13^. Second, the cells used in this experiment were derived from a single cell through clonal expansion ensuring genetic identity. It is well-established that genetically identical mES cells cultured in 2i+LIF have remarkably similar chromatin states and transcriptomes^14,15^. Third and finally, we used fluorescence activated cell sorting to specifically select cells in G2/M phase of the cell cycle, thus excluding major cell cycle effects. This particular combination of growth medium (2i+LIF) and cell cycle phase (G2/M) has been reported to generate particularly homogeneous cell populations with regard to their transcriptome signatures^9^. Using published marker genes, we confirmed that our cells were in the correct cell cycle phase^13,16^ and expressed pluripotency but not differentiation marker genes^9,17^ (Suppl. Fig. 5). Altogether our cells comprise a homogenous population, unconfounded by cell cycle or differentiation effects.

### Discovery of >80,000 significant positive and negative gene covariances

To study pairwise gene covariances, we calculated Spearman’s rank correlation coefficient for all possible gene pairs. We chose this procedure for its ability to detect non-linear monotonous dependencies and for its robustness towards outliers. The calculation of covariance coefficients with Spearman’s was performed separately on each of the three biological replicates. The measured covariance coefficients were centered around zero (Figure 1A, left), indicating the absence of overall confounding factors, and importantly, the observed covariance values had a bigger spread that those of permutated controls (Figure 1A), suggesting the presence of numerous non-random biological covariances. We then defined covariances to be overall significant if their coefficients were significant (p < 0.01) in two out of the three replicates and had the same sign in all three replicates. This approach resulted in 81,820 significantly covarying unique gene pairs (52,695 positively and 29,125 negatively). Using stringent permuted controls, we estimated the false discovery rate of these covariances to be lower than 1.6% (Methods). An example for a highly significant gene pair is shown in Fig. 1B, where each data point represents one individual cell. Significant covariances identified with Spearman’s ranked correlation coefficient have a high overlap with those retrieved by Pearson’s correlation coefficient, and with dependency measures recovered through Hoeffding’s D statistics (Fig. 1C), showing the robustness of the approach. Finally, we validated several of the gene expression covariances using single-molecule FISH and single-cell quantitative RT-PCR (Suppl. Fig. 11, 12). In summary, we present >80,000 high-confidence gene pair covariances; more than have been reported in previous single-cell studies.

**Figure 1:**
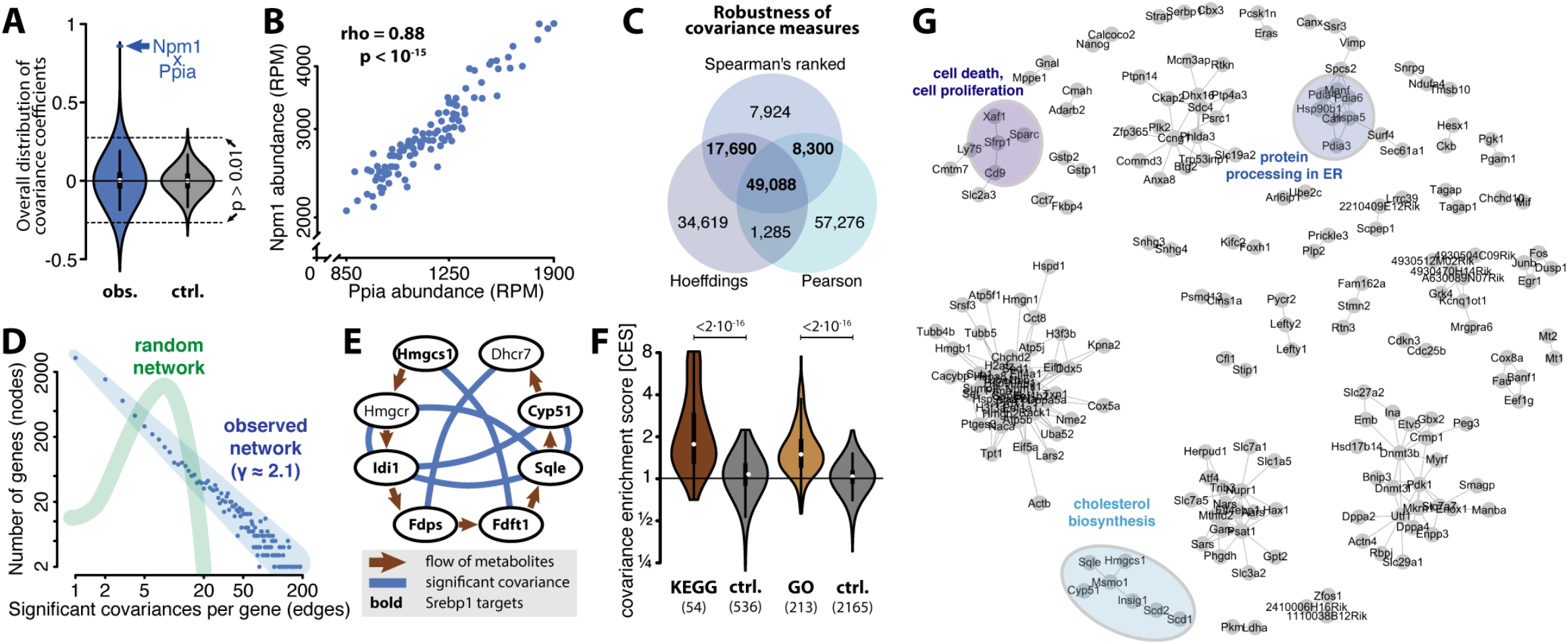
Covariance network reflects biological features. **(A)** Transcriptome-wide covariance (co-expression) values for all possible gene pairs. Violin plot of Spearman’s ranked statistics (rho-value) for the entire transcriptome (blue) and for a permuted control matrix (grey). Value for the gene pair *Npm1* - *Ppia* is highlighted. **(B)** Covariance of the genes *Ppia* and *Npml.* Abundances for the two genes are in reads per million (RPM) and plotted in log scale. Each data point represents their respective measurement in the same single cell. **(C)** Spearman’s ranked coefficients is in accordance with other covariances and dependency measures. **(D)** Gene covariance network is scale-free. Number of significant covariances per gene against the number of genes with that number of covariances. Blue line illustrates a scale-free network that captures the data points (y ≈ 2.1). Green line illustrates the degree distribution of a random network with same number of genes (nodes) and covariances (edges) as the observed network. **(E)** Cholesterol biosynthesis pathway is highly enriched for gene pair covariances. Genes involved in cholesterol biosynthesis from acetyl-CoA. Only genes that were robustly detected in our sequencing data are shown. Arrows indicate the flow of metabolites, lines indicated significant covariance between genes. Gene names in bold indicated direct targets of Srebpf1, a transcription factor that is well known to regulate cholesterol biosynthesis. **(F)** Gene sets that share functional annotations are enriched for covariances. Gene covariance enrichment scores (CES) for gene sets sharing the same gene ontology or sharing the same KEGG pathway annotation as well as respective controls. Gene covariance enrichment scores indicate the ratio of observed significant covariances relative to the amount of expected covariances (see main text). **(G)** Example sub-network.

### Properties of the covariance network reflect biological functions

We observed that the covarying gene pairs link together in complex patterns that can be described as a network. It is well-established that biological networks, such as those arising from transcription factor or protein interactions, have properties that are different from random networks^18^. For instance, biological networks tend to follow power law distributions, such that most genes have only few interactions with other genes while few genes represent hubs in the network, interacting with many other genes. Consistent with our network having biological rather than technical origins, we found that our covariance network follows such a power distribution (γ≈2.1, Figure 1D, light blue). Importantly, this network structure is distinctly different from that of a random network with the same overall connectivity (Figure 1D, light green). Further network features are listed in Suppl. Fig. 7D.

Within the covariance network, we identified many biologically meaningful subnetworks, such as the one formed by genes involved in cholesterol biosynthesis (Fig. 1E). Genes involved in cholesterol biosynthesis are activated when the SREBF1 transcription factor is cleaved from the cell membranes and shuttles to the nucleus in response to lack of cholesterol^19^ and can therefore be expected to covary in expression, depending on the localization of SREBF1 protein. Another notable sub-network is formed by genes involved in the formation of the TCP1 ring complex, a chaperone involved in tubulin biogenesis^20^ (Suppl. Fig. 8).

A substantial part of the observed covariances (~14,500 gene pairs) are between ribosomal proteins. While these are interesting and have been reported before for bulk cell populations^21^, the exact mechanism behind ribosomal protein co-expression in eukaryotes remains cryptic, and we have excluded them from analysis in the sections below. It was recently reported that four of these proteins (RPL10, RPL38, RPS7, RPS25) are optional components of the ribosome, whose inclusion or exclusion can influence which pools of transcripts are preferentially translated^22^. We find that these four ribosomal proteins all covary positively and significantly with each other, providing evidence that they do not function in a mutually exclusive “switch-like” manner in single cells.

Importantly, applying our new method for measuring covariance enrichment over large gene sets (see section of CES score below), we find that genes sharing common Gene Ontology terms are 31% more likely to be covarying (1.31-fold covariance enrichment), while permuted control sets shows no such enrichment (Figure 1F). The same holds true for genes sharing common KEGG pathway annotation, where the enrichment is 25% (Figure 1F). We conclude that genes sharing functions or pathways are more likely to be regulated in a similar fashion and hence tend to covary. In conclusion, the covarying gene pairs form a comprehensive scale-free network, which is associated with annotated cellular functions and pathways.

### Covariances retrace known aspects of stem cell biology

The pluripotency of mouse embryonic stem cells has been studied extensively and several studies focus on characterizing their transcriptomes and gene regulatory circuits^13,23–26^. The network that we observe recapitulates many known relationships between pluripotency markers in mouse embryonic stem cells. For instance, positive covariances support the activation of *Fgf4* through *Nanog* and *Sox2*^27,28^, while negative covariances support the inhibition of *Dnmt3a/b/l* by *Prdm14*^17,29^ and of *Dppa3* by *Tbx3*^30^. While our data support previous claims that Nanog is positively covarying with *Klf4, Sox2, Tet2,* and *Kat6b*^9^, we see little support for a covariance with *Esrrb, Zfp42* and *Tet1* and we observe a significant negative covariance with *Pou5f1* and *Dnmt3a* in single cells. With regard to predicted pluripotency genes, we can confirm that there are strong covariances between *Etv5,* and weak covariances between *Ptma,* and *Zfp710* and other pluripotency genes. Covariances of pluripotency genes can be found in Suppl. Table 4. In summary, the detected covariances are in accordance with known gene expression patterns in stem cell biology, and gives hints to new connections.

### Covariance enrichment score (CES)

To systematically investigate the regulatory implications of the covariances, we defined the *Covariance Enrichment Score* (CES) for gene sets of interest. The CES indicates whether for a given gene set we observe fewer or more significant covariances between the genes than we would expect, based on a simple background model. Our background model considers the total number of significant covariances for each gene as well as the number of covariances of all its potential pairing genes. In short, it is the factor of the probabilities of two genes assuming that their covariances were distributed randomly, summing over all possible pairs in the gene set.

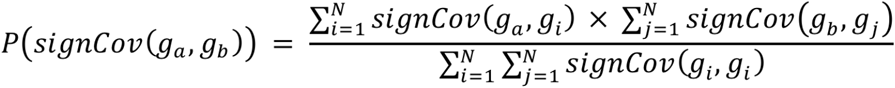

For instance, we assume that genes that are regulated by the same transcription factor will tend to covary, depending on the abundance and activity of the transcription factor in individual cells. In the next sections we test such hypotheses using CES, and in the case of one type of regulator (miRNAs) use a loss-of-function mutant cell line to validate that the observed covariances are indeed a direct effect of said regulator.

### MiRNAs induce transcriptome-wide gene expression covariances

We first apply the Covariance Enrichment Score to study the regulatory impact of miRNAs, which are important post-transcriptional regulators of gene expression (reviewed in Bartel, 2018). In most conditions, these small RNAs down-regulate the expression of protein coding genes by binding their mRNA transcripts and leading to their degradation^32^. This targeting takes place in the cytoplasm and is therefore spatially decoupled from transcriptional regulation.

We speculate that miRNA regulation of gene expression may be a source of gene covariances. For instance, if a miRNA is highly abundant in a given cell, its targets may be coordinately repressed, and we expect an enrichment of covariances for these targets. To test this hypothesis, we investigated the top-ranking miRNA targets according to TargetScan^33^, which is the most widely used catalog of miRNA-target interactions. In this study we focused on the seven highest expressed conserved miRNA families (including the miR-15 and miR-290 families) in mouse embryonic stem cells (Suppl. Table 5).

Strikingly, miRNA gene target sets are significantly enriched for gene covariances. In median the top 200 targets of each of the seven miRNA families are 31% more likely to covary with each other than expected (p=0.032). The enrichments exhibited a gradient with the enrichment being stronger for the top-ranking targets compared to sets that also included lower ranking targets (Figure 2A). Introns are spliced out in the nucleus, so their abundances cannot be impacted by miRNA action in the cytosol. Consistent with this, miRNA targets do not significantly covary at the intron level (Figure 2B).

**Figure 2:**
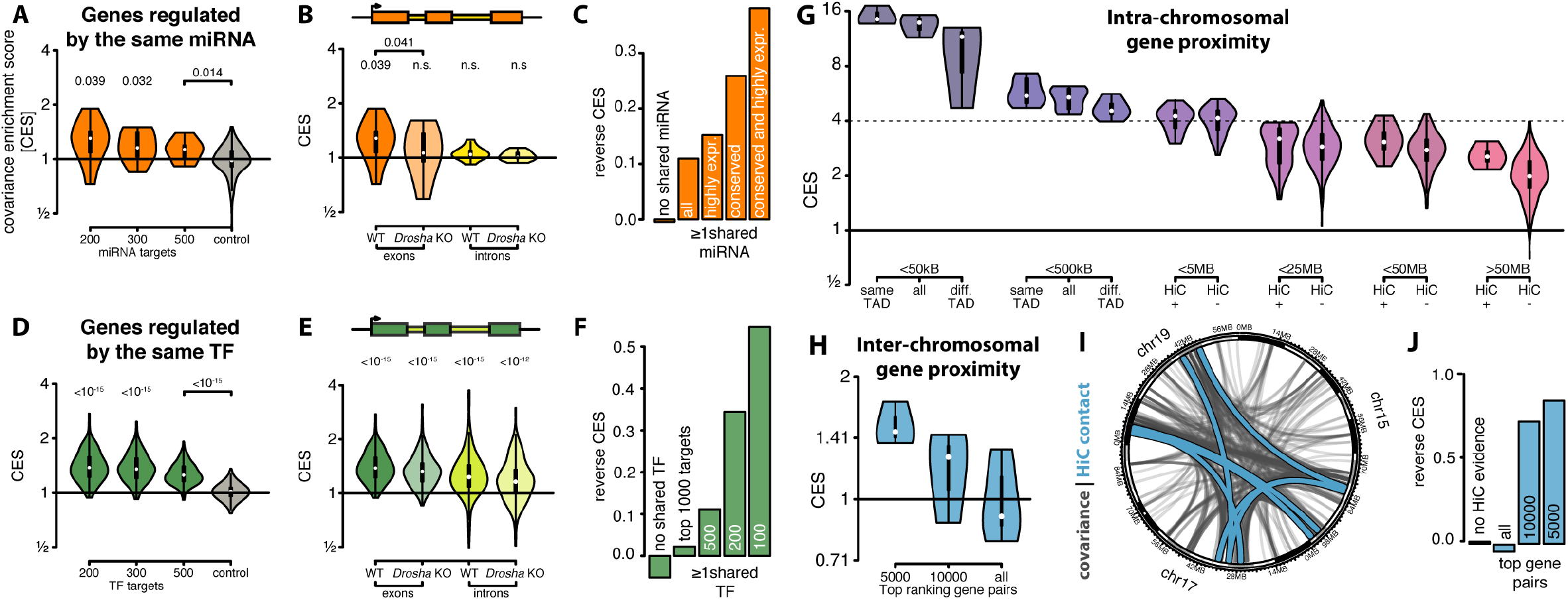
miRNAs, transcription factors and nuclear organization define covariances. **(A)** miRNA targets tend to covary. Covariances enrichment scores (CES) for the top 200, 300 and 500 ranked miRNA targets according to TargetScan, for the seven highest expressed conserved miRNA and for a control set for comparison for 300 randomly selected targets. P-values refer to respective controls. **(B)** miRNA target covariances occur post-transcriptionally and are miRNA-dependent. Enrichment in sets of the top 200 ranked miRNA targets in parental cells (WT) and *Drosha* KO cells, that are void of canonical miRNAs. Enrichments are color coded for exonic reads, representing post-transcriptional regulation (orange) or intronic reads representing transcriptional regulation (yellow). **(C)** Covarying genes are enriched for shared miRNA targeting. Reverse covariance enrichment shows the ratio between covariances that share a common miRNA and permuted covariances that share a common miRNA. **(D)** Transcription factor targets are enriched for gene covariances. Enrichment in sets of the top 200, 300 and 500 transcription factor targets, for 168 transcription factors profiled with ChIP-seq. Control for comparison is shown for 500 randomly selected targets. P-values refer to respective controls. **(E)** Transcription factor target covariances are transcriptional and miRNA-independent. Enrichment in sets of the 200 ranked transcription factor targets in parental cells (WT) and *Drosha* KO cells. Enrichments are color coded for exonic reads (dark green) or intronic reads (light green). **(F)** Covarying genes are enriched for shared transcription factor targeting (figure similar to 2C). **(G)** Genes that are in close nuclear proximity and locate to the same chromosome are enriched for covariances. The range categories are mutually exclusive, for instance pairs of genes that are <5MB apart are not included in the <25MB category. **(H)** Gene regions that are in close nuclear proximity and locate to different chromosomes are enriched for covariances. Since relatively few intra-chromosomal Hi-C contacts were identified, we here used a less stringent criteria (p-value <0.05) cut-off to robustly identify significant covariances (Suppl. Methods) **(I)** Circos plot showing significant covariances and Hi-C contacts for chromosomes 15, 17, and 19. Significantly covarying gene pairs are connected by a light blue line. Inter-chromosomal Hi-C contacts are shown as grey lines. **(J)** Covarying genes are enriched for inter-chromosomal Hi-C contacts (figure similar to 2C).

To exclude the possibility that these covariances originate from other post-transcriptional effectors, we investigated cells that are void of canonical miRNAs. DROSHA is an endonuclease involved in the biogenesis of miRNAs, without which canonical miRNAs cannot be produced. We used an inducible *Drosha* knock-out cell line to validate the miRNA dependence of these covariances (Methods), and sequenced the transcriptomes of 343 of these knock-out cells using clonal expansion from a single cell and sorting of cells in G2/M phase as described above. We have previously demonstrated the global loss of miRNAs in this particular cell line^34^. As expected, there is no covariance enrichment in miRNA target sets in *Drosha* knock-out cells (Figure 2B), demonstrating that these covariances are directly caused by miRNA activity.

We additionally investigated the “reverse covariance enrichment”. Here, we observe the set of all significantly covarying gene pairs and ask how often they are regulated by the same miRNA, compared to a background set (Methods). We find that covarying genes are 12% more likely to be co-regulated by the top 16 miRNAs, and 35% more likely to be regulated by the seven most highly expressed conserved miRNAs (Figure 2C), showing the importance of miRNA conservation and abundance in inducing covariances. It has previously been reported that individual miRNAs can induce gene covariances^35^, but here we for the first time show that this holds for the larger complement of miRNAs, transcriptome-wide, and that natural (non-induced) fluctuations of miRNA abundance or activity are sufficient to cause the covariances.

From a network perspective, we found that >6,000 high-confidence gene covariances were lost in the cells void of miRNAs, while less than 3,000 new covariances were gained (Suppl. Figure 7A). A substantial number of the genes that cease to covary were miRNA targets and the ratio of lost to gained covariances increase when high-confidence targets were considered (Suppl. Figure 7B). The genes that lost covariances were enriched in functions in RNA biology (Suppl. Figure 7F), including regulation of PolII regulation. The average number of covariances per gene decreased significantly in the miRNA-depleted cells, from 10.1 to 8.4 covariances, and the number of genes without covariances increased from 2,265 to 2,866 (Suppl. Figure 7D). Overall, this indicates a global loss of gene expression coordination in cells that are void of miRNAs.

### Genes regulated by the same transcription factors covary with each other

To investigate how regulation by transcription factors influences covariance patterns, we studied the binding sites of 145 transcription factors for which mouse ES cell ChIP-seq data were deposited in the Cistrome database^36^. As for miRNAs, we observe a gradient in covariance enrichment which is stronger for the top-ranking transcription factor targets compared to lower ranking targets (Figure 2D). Importantly, transcription factor target sets are significantly enriched for gene covariances both on the exon and the intron level (Figure 2E), consistent with transcriptional regulation. In median, the top 150 ranked targets of these transcription factors are 39% more likely to covary on the exon level (p-value<10^-15^) and 22% more likely to covary at the intron level (p-value<10^-15^) than are background genes. As expected, the covariance enrichment of transcription factor targets is not significantly lowered in *Drosha* knock-out cells (Figure 2E). In conclusion, genes that are regulated by the same transcription factor tend to covary, possibly due to stochastic variations in transcription factor abundance and activity between individual cells. This effect acts on millions of gene pairs and the mean magnitude of the regulation is similar to what we describe for miRNA-specific regulation.

### Genes in nuclear proximity on same or different chromosomes covary

Genes that neighbor on the same chromosome are known to show co-expression^37^; this also holds for genes within the same chromatin loop or within the same topologically associated domain (TAD). Further, the concept of transcription factories covers dynamically assembled complexes that facilitate transcription and are dependent on intra- or inter-chromosomal interactions^38^. To investigate covariance enrichment on genomic regions that are in proximity within the nucleus, we analyzed mouse embryonic stem cell Hi-C-seq data^39,40^. In the following, we define “proximal” genes as those whose interaction is supported by Hi-C data, whether the interaction is intra- or inter-chromosomal (Methods). Our data shows that genes which are proximal and located on the same chromosome are highly enriched for covariances (Figure 2G). Genes that are close in linear distance on the chromosome (<5 MB) are enriched 4-fold in covariances, while genes that are distal (>50 MB) are enriched 2.1-fold. This effect is also detectable at the intron level, confirming an origin in transcriptional regulation at the level of nascent transcripts (Suppl. Fig. 10). Genes that are on the same chromosome are almost twice (1.9-fold) more likely to covary than expected even when their proximity is not supported by Hi-C (Figure 2G, furthest right). However, the highest enrichment was detected for genes that are in close in linear distance on the same chromosome and predicted to be in the same TAD, which are ~15-fold more likely to covary (Figure 2G, furthest left). Intriguingly, proximal genes on different chromosomes also show substantial covariance enrichment (Figure 2H-J), supporting the notion of transcription factories that incorporate areas from multiple chromosomes.

Next, we ranked the relative importance of transcription factors, miRNAs and nuclear proximity for the regulation of covariation (Figure 3). We found that miRNA targets were 35% more likely to covary, transcription factor targets were 39% more likely to covary, genes in nuclear proximity on different chromosome were 3-fold more likely to covary, and, remarkably, genes that are proximal on the same chromosome are 5.3-fold more likely to covary. In summary, we find that transcriptional regulation, miRNA-mediated regulation, and surprisingly, inter-chromosomal nuclear proximity all play important roles, while the intra-chromosomal nuclear proximity is the strongest predictor of gene expression covariances.

**Figure 3:**
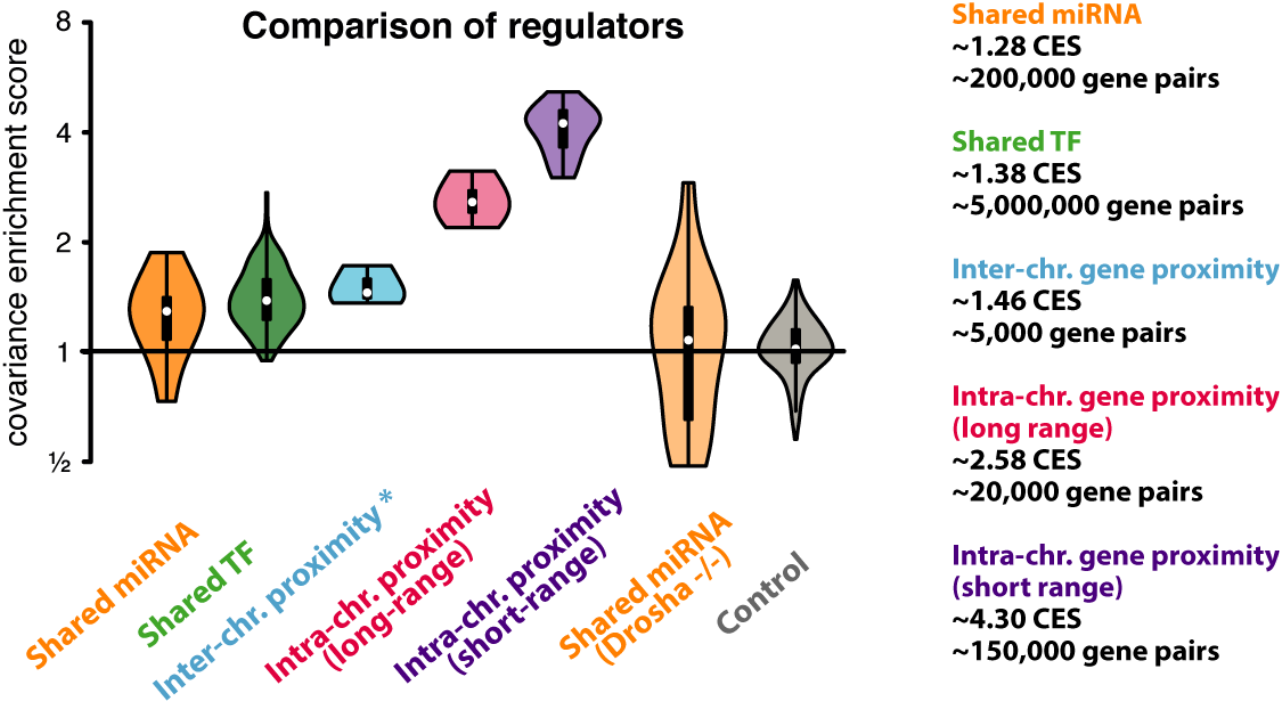
Relative importance of miRNAs, transcription factors and nuclear proximity for covariances. Comparison of covariance enrichment (CES) scores for genes that are either regulated by the same miRNA, regulated by the same transcription factor or that are in nuclear proximity – divided into intra- and inter-chromosomal pairs. The *Drosha* -/- cells are devoid of miRNAs, and the control gene sets are generated by computationally selecting background genes.

### Protein interaction rather than shared function drives gene covariances

Next, we examined putative functions of the covariances that we observe in single cells. We formulated two hypotheses. The first hypothesis can be described as the *“pathway hypothesis”* – that genes involved in the same pathway are coordinated in expression, for instance to avoid bottlenecks in the production of metabolic intermediates^41^. The second hypothesis is the *“complex hypothesis”* – that covariances ensure correct stoichiometry among proteins that are part of the same heteromeric protein complex, since surplus proteins may mis-fold or even cause aggregates^42^.

As stated earlier, genes that share the same Gene Ontology function or process or the same KEGG pathway annotation are significantly enriched for gene covariances (Figure 1F). The same is true for genes that physically interact on the protein level according to experimental evidence gathered by the STRING database (Figure 4A). For these interactions we can see a gradient with those interactions with the highest confidence/affinity score also having the highest enrichment for covariances. We further find that genes that contribute to the same complex are 2.6-fold more likely to covary, compared to just 1.6-fold for genes that are part of the same pathway (Figure 4B), lending support to the “complex hypothesis”.

**Figure 4:**
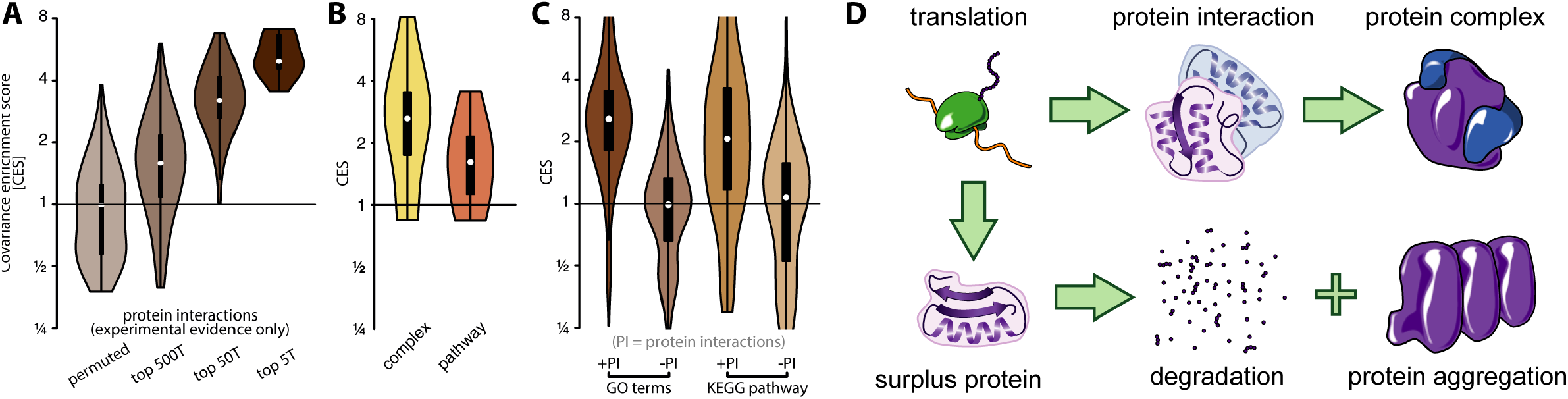
Proteins that physically interact are specifically enriched in covariances at the RNA level. **(A)** Genes that interact on the protein level are highly enriched for covariances. Covariance enrichment of genes that are annotated to be interacting on the protein level according to the STRING database. **(B)** Covariances enrichment for genes whose protein products are part of the same physical complex and for genes that are part of the same reaction (pathway). **(C)** Gene covariances are mainly driven by protein interaction. Covariance enrichment of genes sharing the same GO annotation or KEGG pathway annotation. GO and KEGG annotations are stratified into pairs with shared annotation and experimentally identified protein interaction (+PI) and shared annotation but lack of experimentally identified protein interaction (-PI). **(D)** Model. Heteromeric protein complexes require proper stoichiometry of protein components. Proteins that are in surplus can be degraded, mis-folded or form aggregates.

To further test the two hypotheses, we split the set of functionally related genes into one set with genes that share a functional annotation and protein interaction and those that share functional annotation but no protein interaction. If the “pathway hypothesis” holds, we would expect both gene sets to covary, since they share functions. If the “complex hypothesis” holds, we would expect only the genes whose proteins physically interact to covary, since the covariances are needed for proper stoichiometry of proteins in the complexes. Strikingly, we find no covariance enrichment for either of the GO and KEGG functional annotations if genes that physically interact at the protein level are excluded from the analysis (Figure 4C). In other words, there is no covariance enrichment for proteins in the same pathway, if there is no evidence that they physically interact. Altogether, we present evidence that support direct interactions between proteins in the same complex (Figure 4B-D) as a selector for covariances, rather than pathway stoichiometry.

### Predictive power of gene covariances

We next investigated if our observed covariances can be used to predict genes that share upstream regulators by a “guilt-by-association” principle. We hypothesize that if a gene of interest covaries with numerous known targets of a transcription factor, it is likely a target of said factor. To test this hypothesis, we noted all genes that had been identified as targets of the transcription factor Ctnnb1 in a mouse ES cell ChIP-seq experiment^43^. This gene is known to regulate cell adhesion and has been linked to various cancer phenotypes^44^. We then ranked all other genes according to how many of the top 100 Ctnnb1 targets they covary with, and observed that the more covariances a gene exhibited, the more likely it is to be bound by Ctnnb1 in a second ChIP-seq experiment^43^ (Figure 5A). In other words, the more significant covariances a gene had with the high confidence targets identified in the first experiment, the more often is was observed among the high confidence targets identified in a second independent experiment. While the predictive power of our method is limited (target probability ~ 25%, Figure 5A) it serves as a proof of principle that single-cell transcriptome data can be used for predicting regulatory relations even in a homogeneous cell population. This approach could be used to make sparse data sets more complete, through “guilt-by-association” with previously identified targets or to identify targets that escape current technologies due to biases. Last, we found that the function of genes could be inferred by surveying functional annotations of covarying genes (Figure 5B, Methods). This may not only aid functional annotation but could also reveal hidden gene functions, so called moonlighting. In conclusion, knowledge of gene covariances across single cells can be used to infer gene function and regulation through associations.

**Figure 5:**
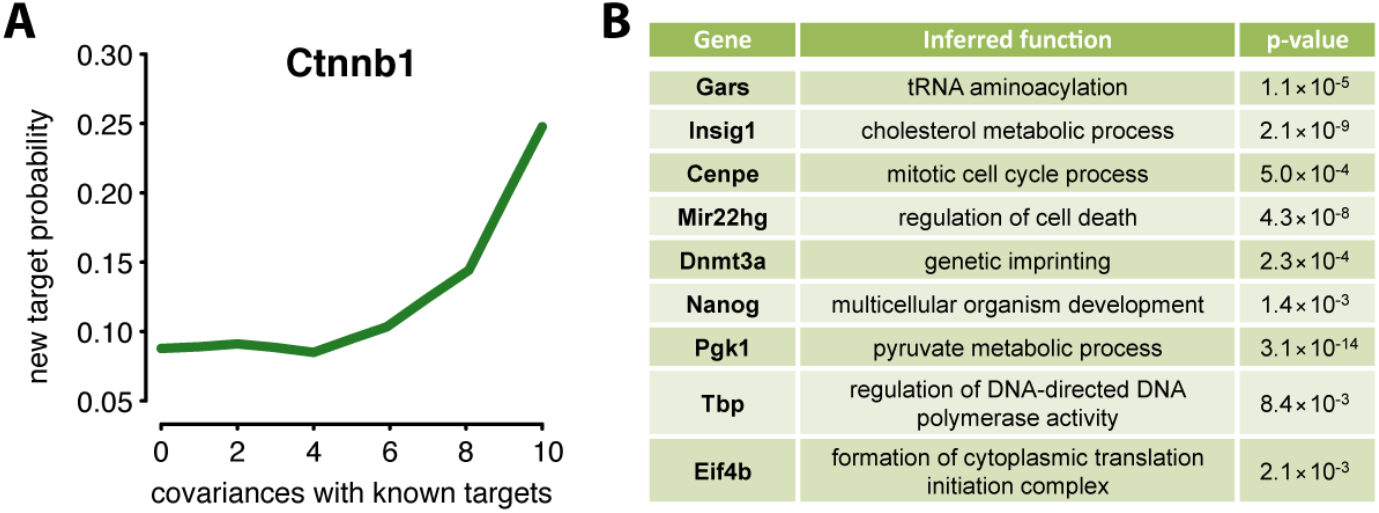
Gene covariance information can predict regulatory targets. **(A)** Genes that covary with transcription factor targets are likely targets of the same factor (Ctnnb1) and can be validated by ChIP-seq. Probability for genes that share a certain number of significant covariances with the top 100 targets identified via ChIP-seq to be identified *de novo* in an independent second ChIP-seq experiment. **(B)** Examples of genes whose function was determined purely from the functional annotation of covarying genes.

## Discussion

We show that statistically robust and biologically meaningful gene covariances can be detected in homogeneous non-dynamic single cell populations. Evidence to support this claim include: the validation by alternative statistical methods, a low estimated false discovery rate, the recovery of known regulatory patterns, and a power-law distribution of network edges commonly found in biological networks. Our experimental set-up allows for the study of widespread gene expression covariances unrelated to cell cycle and other dynamic changes in the cells such as differentiation. Strikingly, all major regulatory mechanisms - post-transcriptional, transcriptional and by nuclear proximity - influence covariance patterns. We confirmed experimentally the importance of post-transcriptional regulation by miRNAs by showing that loss of miRNAs results in a specific loss of a subset of covariances.

Based on our findings, we propose a hierarchy of gene covariance regulation. We place regulation via intra-chromosomal proximity first because of the strength of the effect, and transcription factors second because of the size of the affected target pool (Figure 3). The influence of inter-chromosomal proximity and miRNA regulation is comparatively smaller although still substantial. As targets of the same regulator tend to covary as well as genes that are part of the same functional units, covariances have been used to predict gene function and regulation. We show that we not only recover known gene functions and transcription factor targeting but, as a proof of principle, also showed the predictive potential for both gene function and regulation.

Importantly, we find that covarying genes only tend to share the same function if their encoded proteins also physically interact, suggesting a role in protein complex stoichiometry. The induction of gene expression covariation could be beneficial to cells as it is well understood that the formation of heteromeric protein complexes is often needed for proper folding and stability of the proteins involved^45^. In bacteria, spatial separation of the translation of such proteins leads to misfolding events^46^. It is conceivable that temporal separation might result in similar effects. The production of misfolded proteins that have to be removed by degradation is costly from an energetic point of view, and accumulation of misfolded protein can have lethal consequences for cells (Figure 4D). Therefore, we suggest that establishing expression covariance of such genes already on the RNA level might be an advantage in evolutionary terms.

In this study we measure RNA rather than protein with the latter being closer to the cellular phenotype. However, when inferring upstream regulation, it may be more informative to measure RNA. Further, many of the interesting and biologically meaningful covariances that we discover may not be detectable at the protein level, even in single cells. For instance, transcript covariances may be important for co-folding, but they may not be visible at the proteome level for proteins that have long half-lives and that are therefore stably expressed. It will be exciting to study covariances at the protein level, when technologies to accurately profile hundreds of proteins in single cells become available.

Gene co-expression studies have been conducted on pools of cells for decades, yielding important insights into covariances and network properties. However, these studies have been limited in their capacity to study changes to network properties following a genetic perturbation. For instance, to study the effects of *Drosha* knockout using pooled cells, it would be necessary to ablate the gene in dozens or hundreds of cell lines in parallel to have the statistical strength to call covariances. In contrast, our study serves as a proof-of-concept that it is possible to delete the gene in a single cell line, and then consider each of hundreds of individual cells as an independent condition, thus obtaining the statistical power to resolve network properties in one experiment. In our study we find that many more covariances are lost than gained in the *Drosha* knockout cells, and we observe a general loss of network connectivity. This highlights the importance of miRNAs in maintaining gene expression synchronicity and global gene network connectivity; an insight that would be difficult to obtain with bulk cell or classical single-gene approaches. In summary, we demonstrate that the combination of single-cell sequencing, gene covariance analysis and genetic perturbations can yield insights into robustness of regulatory networks with unprecedented ease and depth.

A previous study of RNA and protein covariances using samples from bulk cell populations^37^ found that neighboring genes on the same chromosomes are often coexpressed at the RNA level, but are however not functionally related and that the covariances do not translate to the protein level. To the contrary, we observe that gene pairs in nuclear proximity that share an interaction on the protein level are in fact 7.5-times enriched compared to background, suggesting a specific co-occurrence of nuclear proximity, RNA covariance and shared function. Further, using a database for bulk cell protein expression covariances^47^, we find that 21% of our observed RNA covariances in fact translate to the protein level, compared to 6% for background genes. The apparent contrast between these results may originate from the fact that the previous study was conducted in immortalized primary cell lines from human individuals^37^, where genetic variants that strongly impact protein levels may have been specifically selected against by evolution. In contrast, temporal fluctuations of protein levels may be tolerated in individual cells from cell lines, allowing more refined measurements. This highlights the advantages of studying variation of gene regulation at the single-cell level.

It has been proposed that while prokaryotes use co-transcribed operons to ensure synchronized expression and stoichiometry of proteins in common pathways or complexes, eukaryotes use post-transcriptional regulation to ensure a similar outcome at the RNA level. The integrated effect of dispersed transcription and coordinated post-transcriptional regulation has been named *‘RNA operons’* or *‘Regulons’*^48^. Our results support that eukaryotic post-transcriptional regulation by for instance miRNAs can coordinate gene expression at the RNA level, but we also provide evidence that substantial functional regulation occurs at the level of nuclear organization, by genes on the same chromosome or by genes that are in proximity although on distinct chromosomes.

## Acknowledgements

We acknowledge the following funding sources: ERC Starting Grant 758397, ‘miRCell’; Swedish Research Council (VR) grant 2015-04611, ‘MapToCleave’; and funding from the Strategic Research Area (SFO) program of the Swedish Research Council through Stockholm University. The computations were performed on resources provided by SNIC through Uppsala Multidisciplinary Center for Advanced Computing Science (UPPMAX). The Smart-Seq2 data were generated by the Eukaryotic Single Cell Genomics (ESCG) and sc-qPCR was facilitated by the Single Cell Proteomics facilities at Science for Life Laboratory, Uppsala. We also thank CRG (the Center for Genomic Regulation) for support in the early pilot phase of the project. Special thanks to Marie Öhman, Johan Elf, and Claes Andréasson, as well as the Kutter and Pelechano labs for their feedback and input.

## Author contributions

J.F., S.C-S., D.B.M, L.Z. and M.R.F. conceived the project. S.D.M. performed sequence mapping, expression quantification and quality control and M.T. performed all further computational analyses, which were supervised by M.R.F. J.F. and S.C-S. performed cell perturbation and sorting experiments. E.G. performed and M.B. supervised smFISH validations. C.J.G. performed single-cell qPCR validations. I.B. contributed to smFISH and single-cell qPCR validations. L.Z. performed and S.O. supervised early pilot phase computational analyses. M.T. and M.R.F. wrote the manuscript, with contributions from all authors.

## Author Information

Sequencing data have been deposited at GEO under accession number XXXX. The authors declare no competing financial interests. Readers are welcome to comment on the online version of the paper. Correspondence and requests for materials should be addressed to M.R.F.

## Methods Summary

*Drosha^KO^* cells were generated using the *Drosha^F^* mouse embryonic stem cell line (Chong et al, 2008) containing the tamoxifen-inducible LoxP - *exon9* - LoxP and a neomycin selection cassette. The *Drosha^KO^* cells were propagated for three passages in 2i media containing mouse LIF. For single-cell sequencing, the *Drosha^KO^* cells were stained with Hoechst-33342 and propidium iodide, then sorted in G2/M using a BD Influx (BD Bioscience). Dual-indexed cDNA libraries were prepared using the Illumina NexteraXT dual index library prep kit following the SmartSeq2 protocol (Picelli: 24056875). cDNA libraries were pooled and 50-bp single-end sequencing was performed on an Illumina HiSeq 2000 platform. The reads were mapped against the mm10 mouse genome using Tophat, and duplicates were removed using Samtools. Only reads that mapped to a unique genome position that overlapped a single gene annotation were considered. Individual cells that did not pass stringent mapping criteria, or that were assigned to S or G1 phase by the cyclone function of the SCRAN package, were discarded. Read counts for each cell was normalized to reads per million (RPM). Genes were subsequently excluded from the analysis if they were not detectably expressed in at least half of all cells in each condition and replicate. Pairs of genes were considered significantly covarying if they were correlated by Spearman’s ranked test (p<0.01) in at least two out of three replicates. Transcription factor targets were retrieved from the Cistrome database and miRNA targets were obtained from TargetScanMouse Release 7.1. Targets were ranked by the total amount of 8mer sites, the total amount of m8-7mer sites and the cumulative weighted context score.

